# Genome Report: Chromosome-scale Genome Assembly of the West Indian fruit fly *Anastrepha obliqua* (Diptera: Tephritidae)

**DOI:** 10.1101/2023.09.22.559041

**Authors:** Sheina B. Sim, Carlos Congrains, Sandra M. Velasco-Cuervo, Renee L. Corpuz, Angela N. Kauwe, Brian Scheffler, Scott M. Geib

## Abstract

The West Indian fruit fly, *Anastrepha obliqua*, is a major pest of mango in Central and South America and attacks more than 60 species of host fruits. To support current genetic and genomic research on *A. obliqua*, we sequenced the genome using high-fidelity (HiFi) long-read sequencing. This resulted in a highly contiguous contig assembly with 90% of the genome in 10 contigs. The contig assembly was placed in a chromosomal context using synteny with a closely related species, *A. ludens*, as both are members of the *A. fraterculus* group. The resulting assembly represents the five autosomes and the X chromosome which represents 95.9% of the genome, and 199 unplaced contigs representing the remaining 4.1%. Orthology analysis across the structural annotation sets of high quality tephritid genomes demonstrates the gene annotations are robust, and identified genes unique to *Anastrepha* species that may help define their pestiferous nature that can be used as a starting point for comparative genomics. This genome assembly represents the first of this species and will serve as a foundation for future genetic and genomic research in support of its management as an agricultural pest.

## Introduction

A major pest of mangoes in Central and South America (Carrillo et al. 2017), *Anastrepha obliqua*, commonly denominated as the West Indian fruit fly, is a major threat to economies that rely on mango production. A true fruit fly in the insect family Tephritidae, order Diptera, *A. obliqua* is one of 34 described species in the *A. fraterculus* group (Norrbom et al. 2012), a polyphagous group of fruit fly pests native to Central and South America. The West Indian fruit fly and related species in the *fraterculus* group cause fruit damage through oviposition into ripening fruit and direct larval damage on fruit flesh to at least 60 fruit hosts (Norrbom 1988). This results in economic consequences from damaged products that cannot be sold at market and, more importantly, loss of market access in regions and economies that are naïve to *Anastrepha*. To control and suppress populations of *A. obliqua* and related fruit fly species, commercial growers employ integrated pest management (IPM) methods that include trapping, the use of chemical pesticides, and field sanitation (Villalobos et al. 2017). tioncurrent to commercial efforts, local, state, and federal departments of agriculture work together to implement the Sterile Insect Technique (SIT) (Vanoye-Eligio et al. 2023; Villalobos et al. 2017) and population genetic and phylogenetic analyses (Barr et al. 2017; tiongrains et al. 2023; Ruiz-Arce et al. 2019) to improve management strategies.

To support pest management efforts, we sequenced and assembled the genome of *A. obliqua* to a chromosome scale from a single male specimen. To achieve this assembly, high-molecular weight (HMW) DNA from a single male wildtype *A. obliqua* was isolated and used for single-molecule, PCR-free library preparation and sequencing. Scaffolding the 15 contigs that made up the five autosomes and X chromosome at a chromosome-scale was achieved through syntenic analysis with the recently assembled genome of the closely related *Anastrepha ludens* (tiongrains et al., unpublished), another member of the *fraterculus* group (Norrbom et al. 2012). When evaluated for completeness, the *A. obliqua genome* assembly achieved greater than 97% of the expected benchmark of universal single copy orthologs (BUSCOs) examined at every applicable hierarchical level available. The methods described provide a framework for producing high-quality and chromosome-scale genome assemblies from single insects.

## Methods & Materials

### Wildtype A. obliqua source

A single male West Indian fruit fly was obtained from the Universidad del Valle in Cali, Colombia. The male specimen was obtained from offspring (F1) reared under laboratory conditions. The parents of the specimens were wild insects obtained from red mombin (*Spondias purpurea*) fruits collected in Chagres (Valle del Cauca, Colombia) (03°07’56.6’’N 76°35’31.4’’W), and mated under laboratory conditions. Mango (*Mangifera indica*) fruits were offered to the parent fruit flies for oviposition in cages (25 × 25 × 25 cm) for 24 h. Then, the fruits were packed in plastic containers containing a layer of vermiculite and topped with organdy fabric, where they remained until the emergence of adults. The F1 adults were kept in cages at 28 ± 3°C, RH of 70 ± 10%, and a photoperiod of 12:12 h L:D. Adults were provided with a solid sugar and yeast diet and a continuous supply of distilled water for 90 days. The male specimen was collected at approximately 30 days of age. The specimen was cold-shocked to immobilize and photograph the taxonomic identification voucher (Figure 1). Following this, the specimen was frozen at - 80°C until further processing.

**Figure 1.**
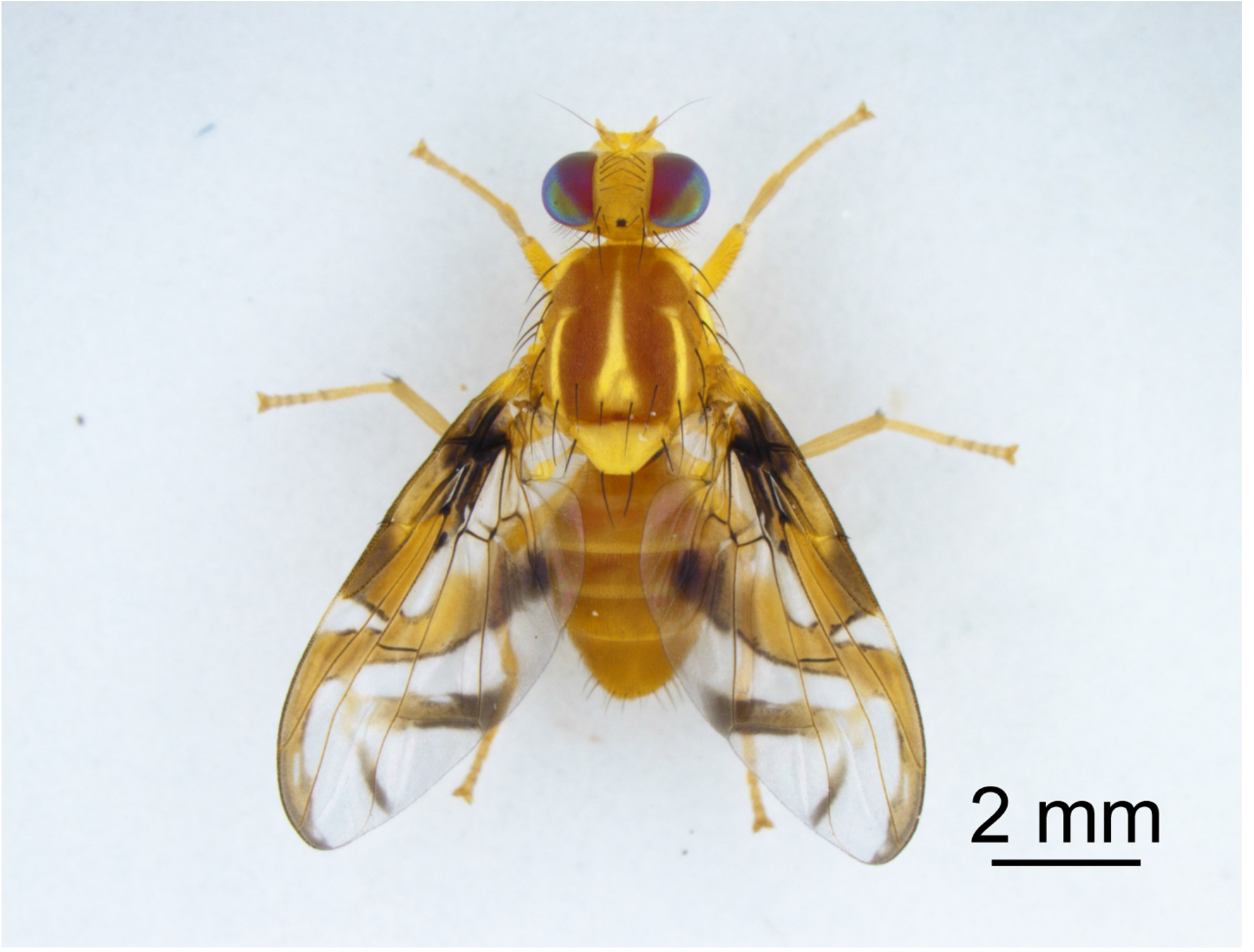
Photo voucher of 30 day old adult male *A. obliqua*. This male *A. obliqua* was the offspring of two wild *A. obliqua* collected from red mombin (*Spondias purpurea*) and ovisposited into mango (*Mangifera indica*). Ayer this photo voucher was taken, this specimen was frozen and homogenized in lysis buffer in preparation for shipment to the USDA-ARS facility in Hilo, HI, USA. Photo credit: Laboratorio Imágenes Postgrado en Ciencias Biología Univalle-Ortega, Velasco-Cuervo.

To prepare the *A. obliqua* sample for shipment from Cali, Colombia to Hawaii, USA, the frozen insect sample was placed in a 1.5 mL DNase-free disposable grinding tube (Thermo Fisher Scientific, Waltham, MA, USA) with 300 µl of Lysis Buffer T1 (Macherey-Nagel, Düren, Germany). The insect was homogenized with a pestle using a vertical motion, avoiding twisting to minimize DNA shearing. The lysate was transferred to a 2.0 mL screw-cap microcentrifuge tube and stored at 4°C until shipment. The sample was shipped on frozen icepacks to the USDA-ARS Daniel K. Inouye US Pacific Basin Agricultural Research Center for further laboratory preparation.

### Library preparation and sequencing

High-molecular weight (HMW) genomic DNA isolation were performed using the MagAttract HMW DNA kit (Qiagen, Hilden Germany) following the protocol for fresh or frozen tissue.

Following isolation, genomic DNA was subjected to a 2.0x bead clean-up to improve sample purity and quantified using the dsDNA Broad Range (BR) Qubit assay (Thermo Fisher Scientific, Waltham, Massachussetts, USA) and the fluorometer of a DS-11 Spectrophotometer and Fluorometer (DeNovix Inc, Wilmington, Delaware, USA). Purity was determined using the UV-Vis spectrometer feature of the DS-11 which reports OD 260/230 and 260/280 ratios. Following the bead clean-up, the HMW DNA sample was sheared to a mean size of 20 kb with the Diagenode Megaruptor 2 (Denville, New Jersey, USA) and resulting size distribution was assessed with the High Sensitivity (HS) Large fragment kit run on the Fragment Analyzer (Agilent Technologies, Santa Clara, California, USA). A PacBio SMRTBell library was prepared using the sheared DNA sample using the SMRTBell Express Template Prep Kit 2.0 (Pacific Biosciences, Menlo Park, California, USA). The prepared library was polymerase bound and sequenced at the USDA-ARS Genetics and Bioinformatics Research Unit in Stoneville, Mississippi, USA on two Pacific Biosciences 8M SMRT Cells on a Sequel IIe system (Pacific Biosciences, Menlo Park, California, USA) beginning with a 2-hour pre-extension followed by a 30-hour movie collection time. After sequencing, circular consensus sequences from the PacBio Sequel IIe subreads were obtained using the SMRTLink v8.0 software.

### Contig assembly

Resulting circular consensus read sequence (CCS) reads were processed by filtering for sequences containing adapter contamination from the raw reads as identified using BLAST+ (Camacho et al. 2009) and a custom database of the PacBio Adapter sequence, C2 primer sequence, and PacBio control sequence following the HiFiAdapterFilt pipeline (Sim et al. 2022). A *de novo* genome assembly using the filtered CCS reads was performed using HiFiASM v 0.11 (Cheng et al. 2021) which results in a primary and alternate assembly containing alternative haplotypes. Both primary and alternate contig assemblies were converted into .fasta format from .gfa format using any2fasta (Seeman 2018). Basic summary statistics on the final primary assembly were calculated using the stats.sh function of BBTools (Bushnell 2014).

### Genome assessment and contaminant removal

Assessment of completeness of the *A. obliqua* contig assembly was determined using the program BUSCO v5.2.2 (Manni et al. 2021; Simao et al. 2015; Waterhouse et al. 2017) and the ortholog database v.10 (odb10) for Diptera. Off-target microbial contigs were identified and filtered using the BlobToolKit pipeline which used nucleotide BLAST (Camacho et al. 2009) and protein DIAMOND BLAST (Buchfink et al. 2021; Buchfink et al. 2015) hits to identify the taxonomic origins of the contigs in the assembly. Adapter contamination was investigated post-assembly as well using the previously described BLAST+ command and custom PacBio adapter contamination database. The contig assembly was initially assessed for the presence of the Diptera BUSCO dataset (odb10) to identify contigs that only contained “Duplicated” BUSCOs and were therefor classified as duplicated contigs that were removed from the primary assembly and included in the alternate assembly. Visualization of the *A. obliqua* genome was performed using the Blobtools2 pipeline (Challis et al. 2020) and a table summary of the Blobtools2 analysis was generated using Blobblurb (Sim 2022). Fidelity of the assembly to the raw data was assessed using YAK which reports a genome QV score (Cheng et al. 2021).

### Mitochondrial genome

All potential copies of the mitochondrial genome were identified using the MitoHiFi pipeline (Uliano-Silva et al. 2023). Briefly, the publicly available mitochondrial genome of *A. fraterculus*, a closely related species, (NCBI GenBank accession KX926433.1) was aligned to the contig assembly using the blastn function of BLAST+ (Camacho et al. 2009). Contigs with high-scoring alignments containing the entire mitochondrial genome were removed from the assembly prior to submission so that it could be manually annotated and submitted separately.

### Synteny analysis

Syntenic relationships between *A. obliqua* and Diptera relatives were used to anchor the *A. obliqua* contigs to Diptera Muller elements and Tephritidae chromosomes. This was achieved by comparing the *A. obliqua* contig assembly to the publicly available chromosome-scale genome assemblies of *A. ludens* (NCBI RefSeq accession GCF_028308465.1) (Congrains et al., unpublished) and *Drosophila melanogaster* (NCBI RefSeq accession GCF_000001215.4) using Diptera odb10 single-copy orthologs. Briefly, *de novo* annotation of the Diptera odb10 single-copy orthologs was performed using MetaEuk (Levy Karin et al. 2020) as part of the BUSCO pipeline (Manni et al. 2021; Simao et al. 2015; Waterhouse et al. 2017) for each of the *A. obliqua, A. ludens*, and *D. melanogaster* assemblies. Following *de novo* annotation, the full table results were joined by gene ID and visualized using the RIdeogram and Tidyverse suite in R (Hao et al. 2020; R Development Core Team 2020; Wickham et al. 2019).

### Scaffolding using synteny

Following synteny analysis, the *A. ludens* genome was used to place 15 *A. obliqua* contigs in a chromosomal context. This species was selected due to its close phylogenetic relationship with *A. obliqua* and because it was assembled to a chromosome-scale. Using the mapped Diptera BUSCOs for both species, contigs in *A. obliqua* were ordered and oriented according to the position of orthologous genes in *A. ludens* using the software ALLMAPS (Tang et al. 2015). Ayer assembly and scaffolding, the final assemblies of *A. obliqua* and *A. ludens* were aligned to each other using Minimap2 (Li 2018; 2021) and visualized using Dot (Nattestad 2017).

### Genome and repeat annotation

Genome annotation was performed using the NCBI eukaryotic genome annotation pipeline (EGAP) and publicly available RNA-seq data from closely related *Anastrepha* species. The genome and resulting annotations from this project currently represent the NCBI RefSeq genome and annotations for *A. obliqua*. Following gene annotation, completeness of the resulting gene set was calculated by performing a BUSCO analysis on the annotations using the ‘-m protein’ option. Annotation of repetitive elements using the EDTA repeat annotation pipeline. Gene annotations and the manually curated transposable elements (MCTE) library for *Drosophila melanogaster (Rech 2021)* were supplied to the EDTA program to mask the genome and improve repeat detection respectively.

### Gene orthology analysis

Gene orthologs for six additional Tephritidae species genomes and one outgroup within Diptera was performed using Orthofinder v2.5.5 (Emms and Kelly 2015; Emms and Kelly 2017; Emms and Kelly 2019). The six species used to determine orthologous genes across Tephritidae were *A. ludens, Ceratitis capitata, Bactrocera oleae, B. dorsalis, B. tryoni*, and *Zeugodacus cucurbitae*. The model species *D. melanogaster* was used as an outgroup (NCBI RefSeq accession numbers for each species can be found in Table S1.). To prepare the protein sequences for every species, the longest isoform per species gene set was first isolated by supplying the RefSeq .gff annotation file to the AGAT function ‘agat_sp_keep_longest_isoform.pl’ (Dainat 2022). This resulted in a gene annotation file for each species in .gff format containing only the longest isoform per gene. Corresponding proteins for each of the longest isoforms was then extracted from the full protein .fasta file for each species and used as the input for OrthoFinder. The OrthoFinder result reporting the number of genes per species for each ortholog group for each species was visualized using the R package UpSetR (Conway et al. 2017) to display numbers of shared orthologs across the datasets.

## Results and Discussion

### Contig and scaffold assemblies

The *A. obliqua* genome was sequenced to 68x coverage and assembled to contigs with a total genome size of 858 Mb (Table 1, Figure 2A). Analysis of completeness using Diptera BUSCOs revealed 97.6% of expected genes found in single copy, 1.5% duplicated, 0.3% fragmented, and 0.6% missing out of the expected 3285 Diptera orthologs (Figure 2A). After placing *A. obliqua* contigs in a chromosomal context and removal of duplicated contigs, mitochondrial contigs, and non-Arthropod contigs as assigned by a nucleotide BLAST search and protein Diamond BLAST search, the final genome size was 837 Mb (Table 1, Figure 2B). Due to the removal of duplicate contigs, the resulting percentage of single-copy orthologs present in the genome increased from 97.6% to 98.1%, and duplicated Diptera BUSCOs decreased from 1.5% to 1.0% in the final scaffolded assembly (Figure 2B). K-mer analysis of the final assembly relative to the PacBio HiFi reads used to create the contig assembly reported a raw quality value (QV) and adjust QV score of 61.388 and 60.790 respectively. This demonstrates that the final genome has high consensus accuracy relative to the sequence data.

**Table 1.**
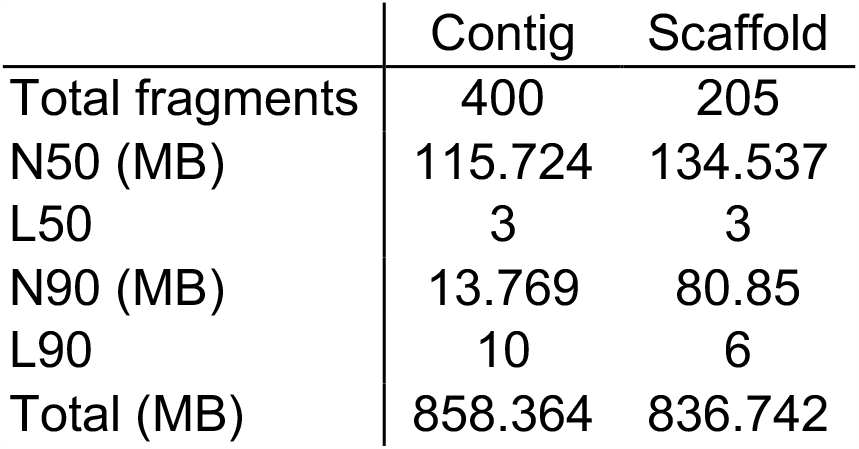
Assembly statistics of the contig and scaffold assemblies.

|  | Contig | Scaffold |
| --- | --- | --- |
| Total fragments | 400 | 205 |
| N50 (MB) | 115.724 | 134.537 |
| L50 | 3 | 3 |
| N90 (MB) | 13.769 | 80.85 |
| L90 | 10 | 6 |
| Total (MB) | 858.364 | 836.742 |

**Figure 2.**
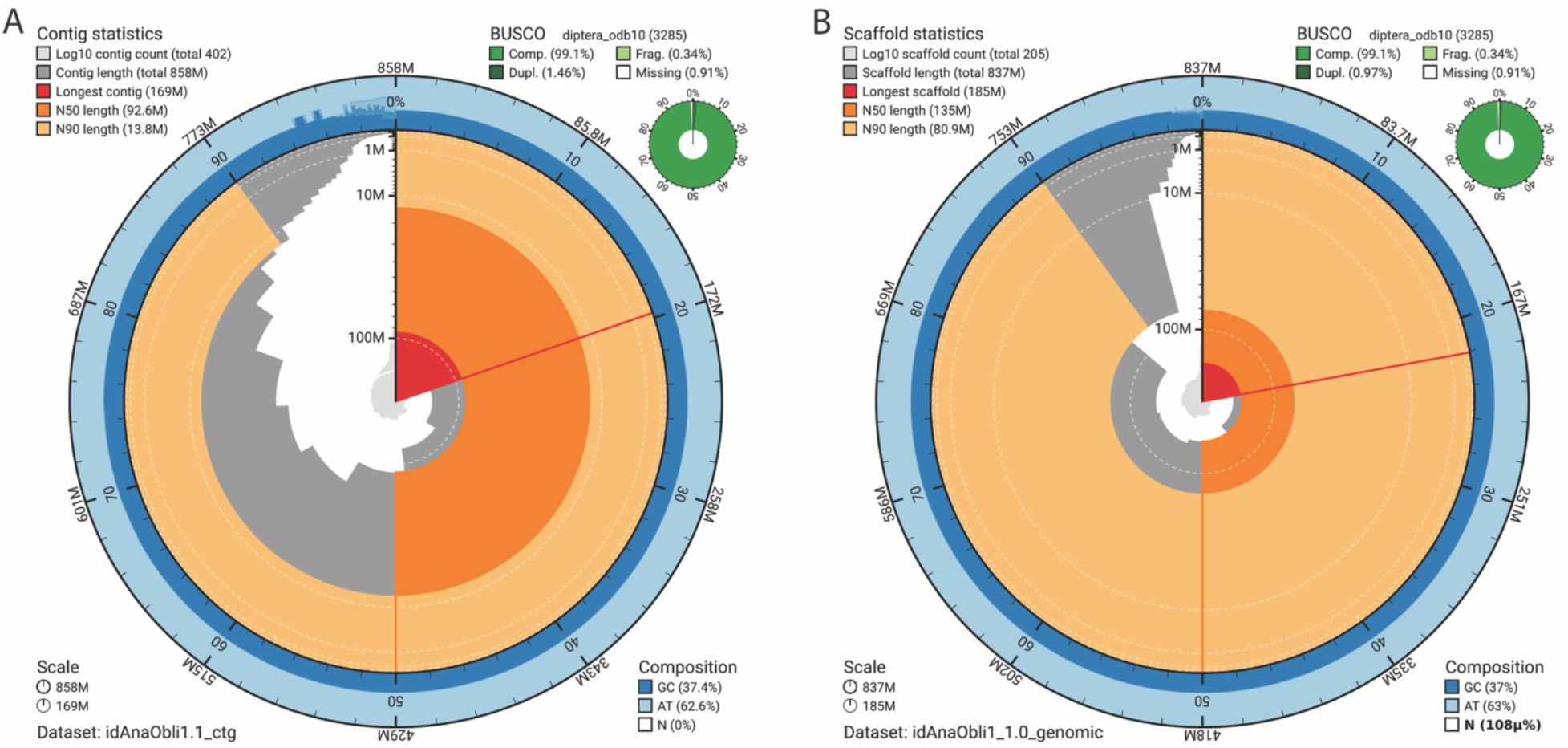
Visual summaries of the contig and scaffolded assemblies. 2A: Snail plot of the contig assembly. The contig assembly output of HiFiASM was in 402 contigs totaling 858 Mb. Depicted are the contigs visually represented as bars around the circle starting at the top and going clockwise and ordered from largest to smallest. In red represents the size and proportion of the longest contig which was reported to be 169 Mb Analysis of completeness as represented by presence of BUSCOs is depicted in the top right of the figure. 2B: Snail plot of the scaffold assembly. After removing non-Arthropod contigs, duplicate contigs, and copies of the mitochondrion, the genome size decreased to 837 Mb, and the assembly was comprised of 6 scaffolds representing the 5 autosome and the X chromosome which represents 96% of the genome, and 199 unplaced contigs and a total of 205 records. BUSCO statistics After duplicate removal showed no change in percentage of complete BUSCOs and a decrease in the number of duplicated BUSCOs to 0.97%. Due to the scaffolding of so few contigs relative to the size of the genome, the proportion of the genome represented by Ns (gaps between contigs) was negligible.

*Ab initio* annotation of the BUSCO Diptera odb10 dataset on the *D. melanogaster, A. ludens* and *A. obliqua* genomes revealed the position of 3187 single-copy orthologous genes in all three assemblies (Figure 3A). Due to high gene collinearity between *A. obliqua* and *A. ludens*, and high synteny in Diptera, synteny analysis enabled the assigning of *A. obliqua* contigs into chromosomes and their corresponding *Drosophila* Muller elements. The single-copy orthologous genes in *A. obliqua* mapped to 15 contigs which represented 95.9% of the genome (802.6 Mb) and anchored the *A. obliqua* contigs into 5 autosomes (Muller elements A-E) and the X chromosome (Muller element F) (Schaeffer et al. 2008). Notably, the *A. obliqua* chromosome that corresponds to Muller element C is completely gapless representing a chromosome-scale contig. Due to low gene density in the X chromosome, orientation of the X chromosome associated contigs requires further validation. Whole-genome alignment of the scaffolded *A. obliqua* and *A. ludens* nucleotide assemblies (Figure 3B) revealed general collinearity across the autosomes with the exception of inversions that were also detected in the gene synteny analysis (Figure 3A). Collinearity across the X chromosome was less than in the autosomes, likely due to the low density of known single-copy orthologous genes across the X chromosome which prohibited the orientation of three of the six contigs assigned to the X chromosome which only had one BUSCO each. Additionally, results from the repeat analysis of the genome revealed that 59.6% of the *A. obliqua* X chromosome was classified as repetitive which is greater than the percent repeat of the entire genome which was 49.46%. This explains the lowered gene content of the X chromosome and poor alignment across the X chromosome.

**Figure 3.**
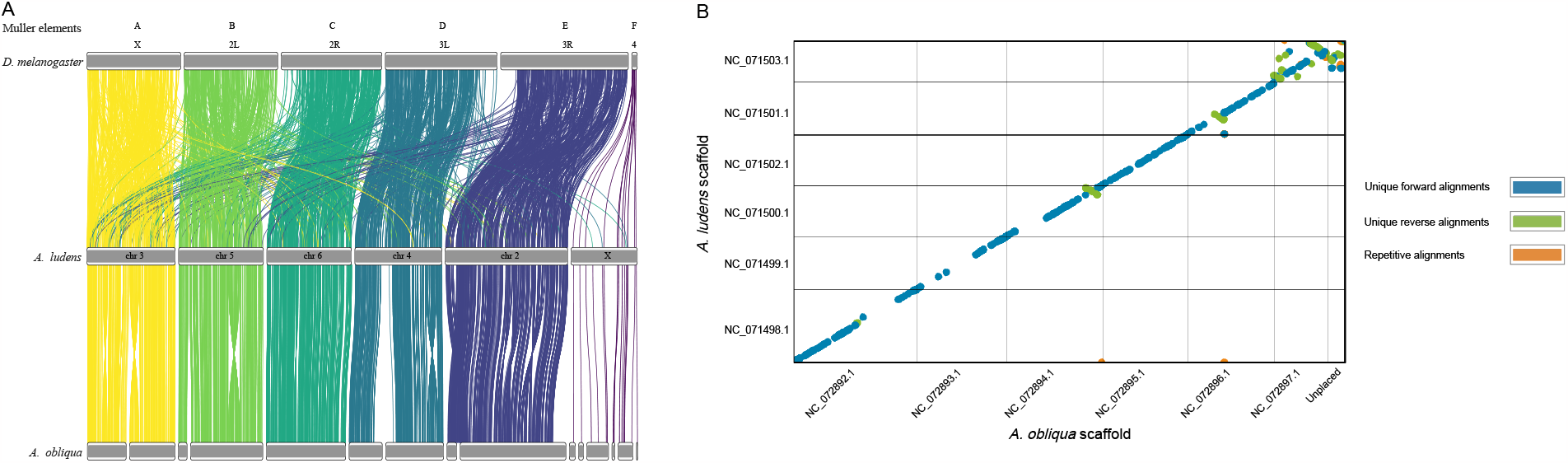
Synteny and whole-genome nucleotide alignment. 3A. The 3187 mapped Diptera BUSCOs assigned *A. ludens* and *A. obliqua* chromosomes to Muller elements in *D. melanogaster*. Each line represents a single BUSCO and its position in the two connecting assemblies (*D. melanogaster* and *A. ludens* on the top half and *A. ludens* and *A. obliqua* on the bottom half). The *D. melanogaster* genome and the *A. ludens* genome have high synteny but low collinearity as there are many genes that are translocated within the corresponding Muller elements and relatively few genes that are translocated between Muller elements. By comparison, the *A. ludens* and *A. obliqua* genomes have both high synteny and high collinearity as no genes translocated between chromosomes were detected in this BUSCO analysis though some chromosomal inversions depicted by the twisted ribbons between the assemblies were observed. 3B. Whole-genome nucleotide alignment between the final *A. obliqua* and *A. ludens* genomes revealed similar collinearity and chromosomal inversions to the synteny analysis. The alignment also revealed that the X chromosome lacked the level of collinearity of the autosomes, likely due to repetitive elements in the X chromosome which lowers the quality of the nucleotide alignment and low gene density in the X chromosome which prohibited the orienting of three of the six contigs assigned to the X.

### Genome and repeat annotations

Genome annotation using the NCBI EGAP detected a total of 14,353 protein-coding genes, 2,033 non-coding genes, 1,820 non-transcribed pseudogenes, 3,622 genes with variants, and no transcribed pseudogenes, immunoglobulin/T-cell receptor gene segments, or other genes outside of the listed categories (Table S2-3). Results of the BUSCO analysis on the gene set showed presence of 99.6% of the expected single copy orthologs in the Diptera odb10 dataset where 97.9% were detected in single copy, 1.7% were duplicated, 0.1% were fragmented, and 0.3% were missing out of 3285 genes. This demonstrates congruence in completeness between the annotated gene set and the genome.

Repeat analysis revealed that 49.46% of the genome contained repetitive elements which represents 413.9 Mb of the 836.7 Mb genome (Table S4). This contrasts with the 39.8% repeat content predicted by GenomeScope2 (Figure S1). This indicates that the difference between the genome size predicted by GenomeScope2 and the final genome assembly size was largely due to underestimation of repetitive elements in the k-mer based analysis of GenomeScope2.

### Gene orthology across Tephritidae

Orthology analysis revealed 15638 orthogroups across all eight datasets which included seven tephritids and one non-tephritid outgroup, *D. melanogaster*. Out of the 15638 orthogroups, 9232 orthologs (59.0%) were found in all the taxa and 7199 were found in single copy in all taxa with the remainder duplicated in one or more species. Though most genes were shared across the Diptera analyzed, the a significant number of genes were unique to Tephritidae (851), unique to *Anastrepha* (360), and 41 were unique to *A. obliqua*. Collectively, these represent novel genes with potential applications to pest management of *A. obliqua, Anastrepha* species, and tephritid species (Figure 4).

**Figure 4.**
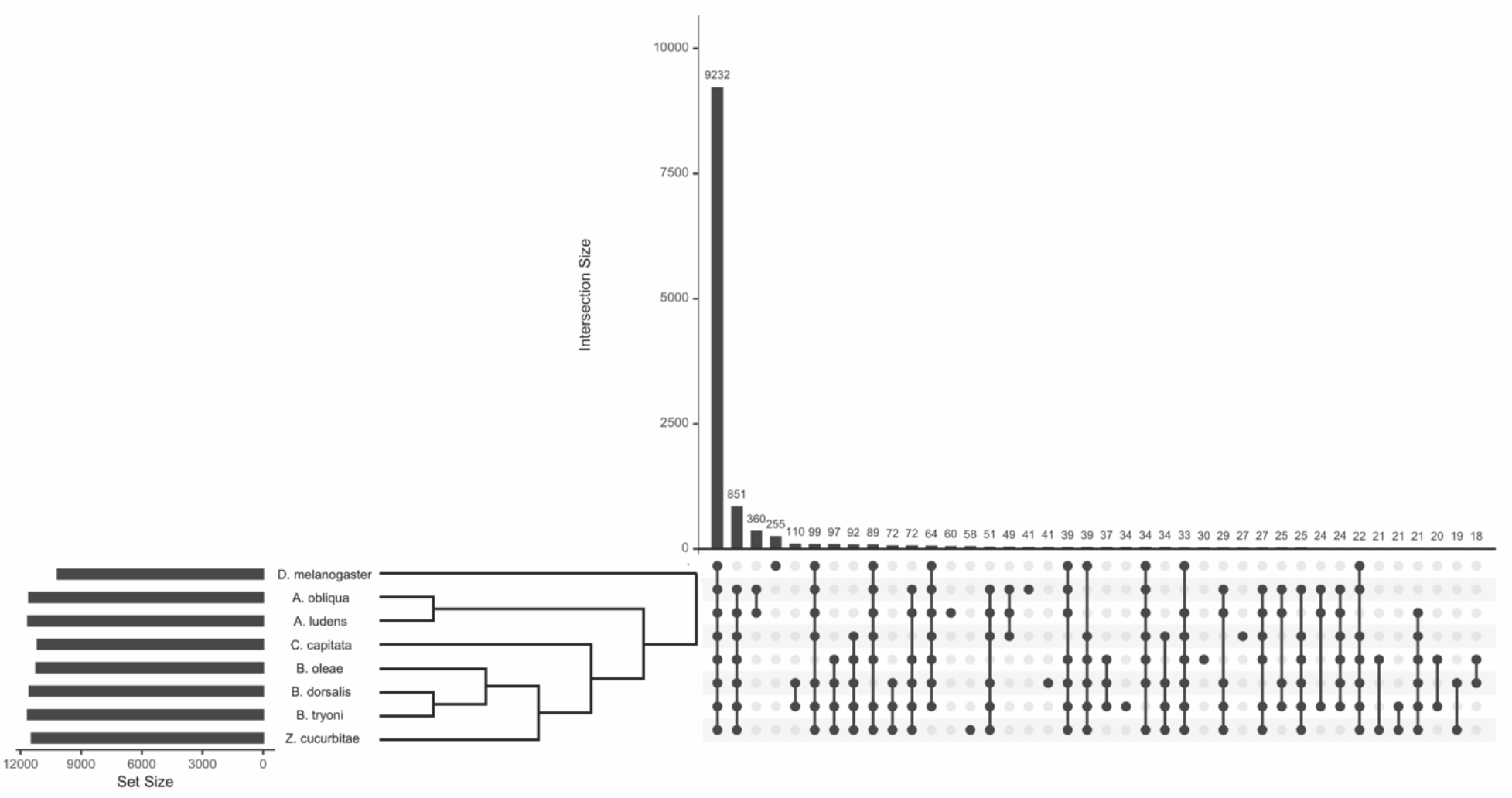
Orthologous genes across Tephritidae and one non-tephritid Diptera outgroup. Orthology analysis of protein sets across *A. obliqua*, six additional tephritid species (*A. ludens, C. capitata, B. oleae, B. dorsalis, B. tryoni*, and *Z. cucurbitae*), and *D. melanogaster* as a non-tephritid Diptera outgroup revealed that the majority of orthologous genes are shared within all the Diptera taxa included in the analysis. Though the majority (59.0%) of genes are shared, there are a large number of orthogroups unique to Tephritidae (851 orthologs, 5.4%), unique to *Anastrepha* (350, 2.3%), and unique to *A. obliqua* (41, 0.3%) that represent novel genes within pestiferous groups that can be investigated for relevance to pest management applications. The phylogenetic relationship between these taxa supports that the most closely related taxa share the most ortholog groups. The number of genes in each dataset is represented in the horizontal bar plot next to the species names and show that the number of genes per species were relatively equal. The numbers above the vertical bars of the upset plot represent the number of genes in that set.

### Conclusions

The reported *A. obliqua* reference genome is the first genome for this species and will serve as a foundation for future pest management methods. It can inform population genetic analyses by anchoring previously identified informative loci and associating them with genes under selection or local adaptation (Aguirre-Ramirez et al. 2021). This high-quality reference genome will also support phylogenomic and species diagnostic analyses by facilitating the identification of loci that can define morphologically cryptic but genetically distinct species (Congrains et al. 2023). Additionally, the annotated gene set can be used in future studies to identify potential targets for genetic control such as RNAi (Maktura et al. 2021) or identifying genes that can be exploited to support *A. obliqua* sterile insect technique SIT programs (Roque-Romero et al. 2022; Roque-Romero et al. 2020). The *A. obliqua* genome represents one of the first two *Anastrepha* species sequenced and assembled at a chromosome-scale and this genome will serve as a genomic resource for future work involving the management of *A. obliqua* and potentially other members in the speciose *A. fraterculus* group.

## Data availability

The primary and alternate assemblies are accessible through the NCBI BioProjects PRJNA796007 and PRJNA797107 respectively. The NCBI RefSeq assembly and annotations are accessible under accession number GCF_027943255.1. The single individual used in this genome assembly is represented under NCBI BioSample ID SAMN24809119. Raw PacBio whole-genome sequences were deposited in the NCBI Sequence Read Archive (SRA) under accession number SRR17536014.

## Acknowledgments

The authors would like to acknowledge D.F. Paulo for valuable contributions to the manuscript, S. Simpson for performing the polymerase binding step and sequencing the sample, and the Laboratorio de Imágenes del Postgrado en Ciencias Biología at the Universidad del Valle and to the biologist Juan Felipe Ortega for his collaboration in photographing and processing the image of *A. obliqua. Anastrepha obliqua* was sequenced as part of the I5K and USDA-ARS Ag100Pest initiative.

## Figures

Figure S1. GenomeScope2 linear plot displaying calculated k-mer abundance and distribution of the adapter filtered PacBio HiFi data used for genome assembly. Analysis of k-mer abundance and distribution using GenomeScope2 revealed that the genome size was estimated at 793 Mb and was sequenced at a depth of about 68x.

## Tables and descriptions

Table S1. Species names and NCBI RefSeq Accession numbers for the protein datasets used in the orthology analysis.

Table S2. Table summary of NCBI Eukaryotic Genome Annotation Pipeline results. Table S3. Table summary of gene annotation characteristics.

Table S4. Table summary of EDTA repeat analysis results.

